# Antibodies control metabolism by regulating insulin homeostasis

**DOI:** 10.1101/2021.08.10.455644

**Authors:** Timm Amendt, Gabriele Allies, Antonella Nicolò, Omar El Ayoubi, Marc Young, Tamás Röszer, Corinna S. Setz, Klaus Warnatz, Hassan Jumaa

## Abstract

Homeostasis of metabolism by hormone production is crucial to maintain physiological integrity and disbalance can cause severe metabolic disorders such as diabetes mellitus. Here, we show that antibodies recognizing insulin are key regulators of blood glucose and metabolism controlling insulin concentrations. In fact, antibody-deficient mice and immunodeficiency patients show sub-physiological blood glucose, which becomes normal after total IgG injection. We show that insulin-specific IgG antibodies found in the serum of wildtype mice or healthy individuals are responsible for this regulation. Interestingly, we identify two fractions of anti-insulin IgM which differ in their affinity to insulin. The low affinity IgM fraction (anti-insulin IgM^low^) neutralizes insulin and leads to increased blood glucose while the high affinity IgM fraction (anti-insulin IgM^high^) protects insulin from neutralization by anti-insulin IgG thereby preventing blood glucose dysregulation. In contrast to anti-insulin IgM^high^, anti-insulin IgM^low^ binds to dsDNA suggesting that it is multi-specific. This multi-specificity mediates the formation of larger immune complexes containing insulin which results in increased uptake and degradation of insulin by macrophages in the presence of anti-insulin IgM^low^ as compared to anti-insulin IgM^high^. To demonstrate that high affinity anti-insulin IgM acts as protector of insulin and counteracts insulin neutralization by anti-insulin IgG, we expressed the variable regions of the same anti-insulin antibody as IgG or IgM. Strikingly, only the anti-insulin IgM regulated insulin function and prevented IgG-mediated neutralization of insulin and subsequent blood glucose dysregulation. Since anti-insulin IgM^high^ is generated in the course of an immune response and affinity maturation, its protective role suggests that preventing autoimmune damage and maintaining physiological homeostasis requires adaptive tolerance mechanisms that generate protective IgM antibodies during memory responses.

## Introduction

Maintaining homeostasis is a complex interplay of the hormone system, immune system and metabolism (Modell *et al*, 2015; Crimeen-Irwin *et al*, 2005). Antibody secreting B cells are fundamentally involved in immunity as they are able to recognize an enormous number of epitopes due to random rearrangement of variable (V), diversity (D) and joining (J) gene segments (Hozumi & Tonegawa, 1976; Roth, 2014). B cells originate from hematopoietic stem cells (HSC) in the bone marrow, where in early stages two IgM heavy chains and two surrogate light chains form the pre-B cell receptor (pre-BCR) (Tsubata & Reth, 1990). Autoreactive specificities are believed to be subjected to removal by clonal deletion or receptor editing (Gay *et al*, 1993; Nemazee & Buerki, 1989; Goodnow *et al*, 1988). Immature B cells expressing an IgM-class B cell antigen receptor (IgM-BCR) leave the bone marrow and home to the spleen where they mature further (Pieper *et al*, 2013). Maturation of B cells in the periphery includes downmodulation of IgM and upregulation of the IgD-class BCR (Übelhart *et al*, 2015; Setz *et al*, 2019; Hartley *et al*, 1991; Roes & Rajewsky, 1993). Recently, our group has shown that the IgD-class BCR is exclusively responsive to polyvalent antigen (Übelhart *et al*, 2015) and that modulation of IgG responses via soluble (auto)-antigens *in vivo* also occurs in an IgD-dependent manner (Amendt *et al*, 2021). Further, mice as well as humans harbor a large number of autoreactive IgD^+^ B cells (Hao *et al*,2011; Cancro, 2020) of which the function remains unclear.

Autoreactive B cells and plasma cells that secrete autoantibodies of the IgG isotype might cause autoimmune diseases such as systemic lupus erythematosus (SLE) (Kaul *et al*, 2016), type 1 diabetes (T1D) (Taplin & Barker, 2008) or rheumatoid arthritis (RA) (Itoh *et al*, 2000; Johnson & Faulk, 1976; Smolen *et al*, 2018). Here, autoreactive IgG is capable of neutralizing its autoantigen or opsonizing it for further degradation or destruction by the innate immune system (Zhang *et al*, 2015). Further, autoantibody-containing immune complexes are able to induce inflammation (Franceschi *et al*, 2017; Suurmond & Diamond, 2015). Another class of autoreactive antibody is resembled by polyreactive natural IgM (nIgM) that has been extensively studied (Boes, 2000; Grönwall *et al*, 2012). nIgM is secreted by B1-B cells and known to clear oxidized lipids and other potentially harmful self-molecules (Tsiantoulas *et al*, 2012). Thus, nIgM is able to suppress sterile inflammation and therefore required to be secreted in an antigen-encounter independent fashion (Boes, 2000; Grönwall *et al*, 2012). In sum, autoreactive IgG and IgM are known to remove self-targets and disease outbreak is dependent on the isotype present.

In mice, we recently identified an autoreactive IgM class that is able to bind and stabilize its cognate antigen. We refer to this class of autoreactive IgM as protective regulatory IgM (PR-IgM) (Amendt & Jumaa, 2021). A key characteristic of PR-IgM is high-affinity and mono-specificity to its antigen. Since PR-IgM is induced in the course of autoreactive immune responses and results in the protection of cognate autoantigen from degradation or uptake. We refer to this phenomenon as adaptive tolerance. However, the presence of PR-IgM in humans is yet unclear.

Here, we show that PR-IgM specific for insulin accumulates with age in human and is able to stabilize Insulin *in vivo*. Furthermore, we show that autoreactive IgG in humans and mice is involved in fine-tuning of metabolic homeostasis by regulating physiological insulin concentrations. In fact, antibody-deficiency is associated with dysglycemia, which is restored after transfer of total IgG from healthy donors.

## Results

### Anti-insulin IgG regulates blood glucose concentration

Using insulin as a common and important autoantigen, we recently developed a simple animal model for testing the induction of autoreactive antibody responses (Amendt & Jumaa, 2021). Importantly, we were able to show that the presence of polyvalent insulin was sufficient to induce an autoimmune response which can easily be monitored by the deregulated glucose concentration in blood and urine similar to diabetes (Katsarou *et al*, 2017; DeFronzo *et al*, 2015). During these experiments we noticed that a considerable amount of total IgG isolated from naïve wildtype (WT) mice was reactive to insulin (Fig. 1A & B). To confirm these data, we performed ELISpot assays and found that anti-insulin IgG secreting B cells are present in the spleen of WT mice (Fig. 1C). When we measured the blood glucose concentrations in WT and B cell-deficient mice, which cannot produce antibodies (Suppl. Fig. 1), we detected a surprising difference. Unexpectedly, the B cell-deficient mice showed abnormally reduced blood glucose levels as compared to WT controls (Fig. 1D).

**Figure 1.**
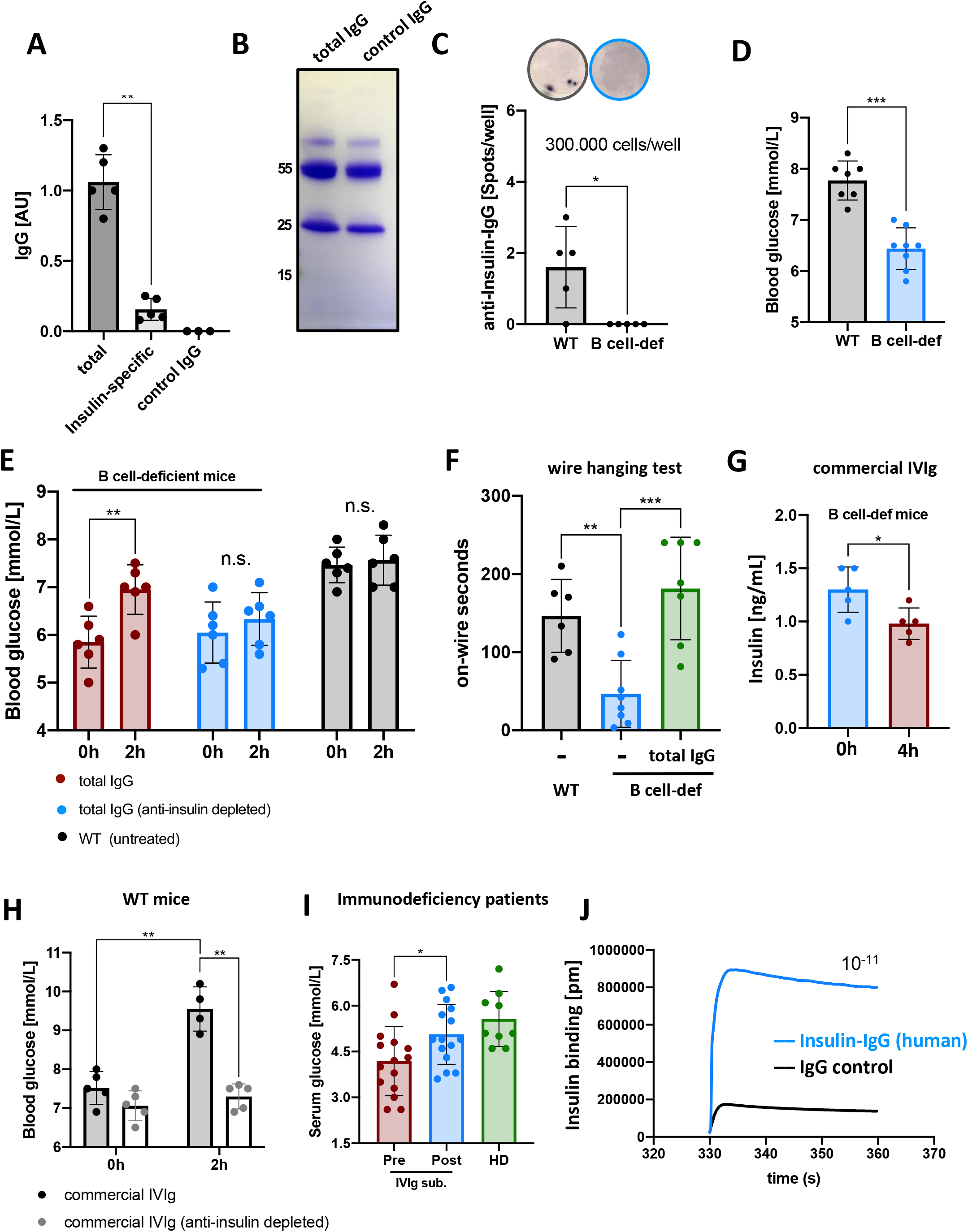
Autoantibodies are required to balance homeostasis in mice. **A:** Insulin-specific IgG concentrations of different IgG pulldowns measured via ELISA (coating: native Insulin). Total: total IgG pulldown via protein G (n=5), Insulin-specific: IgG pulldown via Insulin bait column (n=5), control IgG (n=3). Mean, ± SD, statistical significance was calculated using Kruskal-Wallis test. AU = μg/mL. **B:** Coomassie-stained SDS-PAGE showing total IgG (pulldown from serum) and IgG control (total IgG depleted for anti-Insulin-IgG). Image is representative of three independent experiments. Marker on the left is shown in kilodaltons (kD). **C:** Anti-Insulin-IgG secreting splenocytes of naïve wildtype and B cell-deficient (B cell-def) mice measured by ELISpot (coating: native Insulin). Cells were seeded at 300.000 cells/well and incubated for 48 hours (n=5/group), mean, ± SD, statistical significance was calculated using Kruskal-Wallis test. Top images of wells are representative of three independent experiments. **D:** Blood glucose levels of naïve wildtype and B cell-deficient mice (B cell-def) measured with a commercial blood glucose monitor (mmol/L). Mean, ± SD, statistical significance was calculated using Kruskal-Wallis test. **E:** Blood glucose levels of wildtype and B cell-deficient (B cell-def) mice (red: n=6, blue: n=6) intravenously injected with 200 μg total IgG, IgG depleted for anti-lnsulin-IgG measured at indicated hours. Mean, ± SD, statistical significance was calculated using repeated measure ANOVA test. **F:** Motor function of wildtype (WT) and B cell-deficient (B cell-def) mice as measured by wire hanging test (on-wire seconds). Grey: WT untreated (n=6), blue: B cell-def untreated (n=8), green: B cell-def injected with 200 μg total IgG (n=7). Mean, ± SD, statistical significance was calculated using Kruskal-Wallis test. **G:** Serum insulin concentrations of B cell-deficient (B cell-def) mice injected with 200 μg commercial human IVIg as measured by ELISA at indicated time points (n=5). Mean, ± SD, statistical significance was calculated using Kruskal-Wallis test. **H:** Blood glucose levels of wildtype mice injected with 200 μg commercial human IVIg (black, n=5 0h, n=4 2h p.i.) and commercial human IVIg depleted for anti-Insulin-IgG (grey, n=5) measured by a commercial blood glucose monitor (mmol/L) at indicated hours. Mean, ± SD, statistical significance was calculated using repeated measure ANOVA test. **I:** Serum glucose levels of immunodeficiency patients (SCID) that received commercial IVIg before (pre, n=15) and after (post, n=15) infusion compared to healthy donors (HD, n=9). Mean, ± SD, statistical significance was calculated using Kruskal-Wallis test. **J:** Insulin-binding affinity of human anti-insulin-IgG determined by bio-layer interferometry (BLI). The K_d_ (dissociation constant) was calculated by using the K_a_ (association constant): 1/K_a_. Data are representative for three independent experiments performed in three dilution steps.

To test whether this abnormal decrease is caused by antibody deficiency, we injected total IgG from WT mice, or an anti-insulin IgG depleted control of the same total IgG, intravenously into B cell-deficient mice. We found that blood glucose concentration increased with the total murine IgG, but not with the anti-insulin IgG depleted control (Fig. 1E). In order to test the consequence of reduced steady-state blood glucose on the fitness, we performed wire hanging tests to assess motor function and found that B cell-deficient mice show significantly reduced wire hanging times as compared to WT controls. Importantly, this deficit in wire hanging times was restored after intravenous injection of total murine IgG (Fig. 1F). In addition, B cell-deficient mice also showed dysregulated blood glucose levels after rotarod exercise (Suppl. Fig. 2).

Since total IgG preparations from healthy donors are often used as intravenous immunoglobulin (IVIg) infusions in the treatment of immunodeficiency (Albin & Cunningham-Rundles, 2014) we tested the presence of anti-insulin IgG in these preparations. All preparations contained substantial amounts of anti-insulin IgG. However, the anti-insulin IgG concentration seemed to be increased if the USA was the country of origin (Suppl. Fig. 3). Since insulin is highly conserved between man and mouse, we injected human IVIg into the B cell-deficient mice and detected a decrease in insulin concentration (Fig. 1G). Moreover, injecting 50 μg of human IVIg into WT mice led to increased blood glucose and this effect required anti-insulin IgG since depletion of the anti-insulin IgG from human IVIg prevented the IVIg-induced increase in blood glucose (Fig. 1H, Suppl. Fig. 4).

To test whether the IVIg injection shows similar results in human patients suffering from antibody deficiency, we monitored blood glucose before and after IVIg injection. Similar to B cell deficient mice, antibody deficient patients showed reduced blood glucose concentrations as compared to healthy donors. Importantly, the concentration of blood glucose increased and reached normal levels briefly after the following IVIg infusion (Fig. 1I). Further, immunodeficiency patients that received IVIg showed decreased serum insulin levels immediately after the infusion (Suppl. Fig. 5).

To show that the anti-insulin IgG present in IVIg is specific for insulin, we determined the affinity via bio-layer interferometry (BLI). A dissociation constant of 10^−11^ suggests that the anti-Insulin-IgG is highly specific for insulin (Fig. 1J).

These data suggest that anti-insulin IgG is present in healthy individuals and might be required for the regulation of blood glucose concentration.

### Blood glucose is regulated by antibody class and affinity

To further confirm our finding regarding the presence of anti-insulin antibodies in healthy individuals, we monitored the level of anti-insulin IgG and IgM in the blood of different age groups. We found that anti-insulin IgG was similar in young and aged humans (Fig. 2A), while anti-insulin IgM seemed to decline with age in males and females (Fig. 2B). Interestingly, the human anti-insulin IgM recognizes multiple epitopes of insulin (Suppl. Fig. 6).

**Figure 2.**
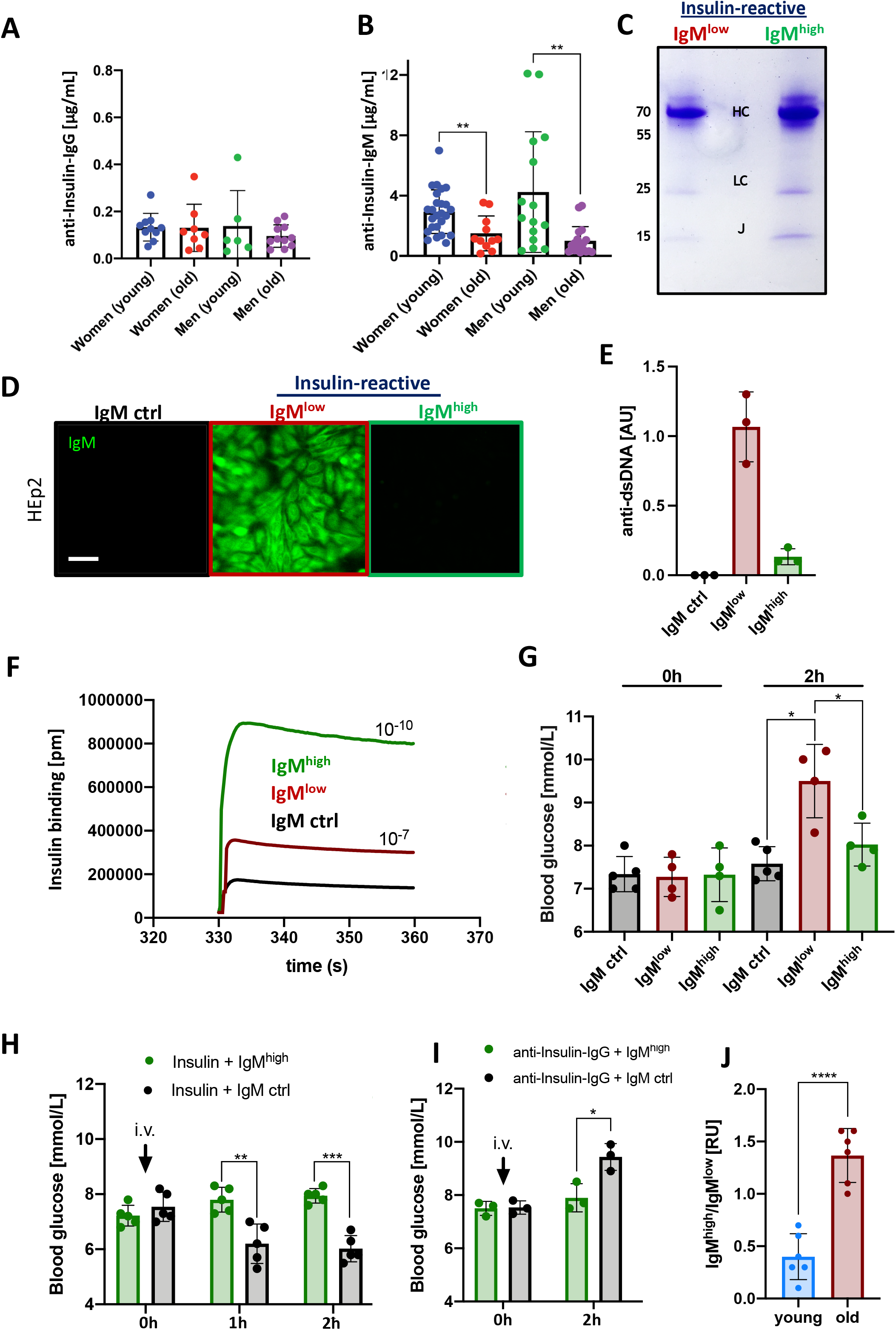
Neutralizing and PR-IgM is present in humans. **A, B:** Serum anti-Insulin-IgG **(A)** and -IgM **(B)** concentrations of young (< 30 years) and old (> 65 years) individuals measured via ELISA (coating: native Insulin). Women (young): n=25, women (old): n=11, men (young): n=15, men (old): n=12. Mean, ± SD, statistical significance was calculated using Kruskal-Wallis-test. **C:** Coomassie-stained SDS-PAGE showing low-affinity anti-Insulin IgM (red) and high-affinity anti-Insulin-IgM (green) after purification. Image is representative of three independent experiments. Marker on the left is shown in kilodaltons (kD), HC (heavy chain): 70 kD, LC (light chain): 25 kD, J (J-segment): 15 kD. **D:** HEp2 slides showing nuclear strucutre-reactive IgM (ANA) of insulin-reactive IgM pulldowns. Black: monoclonal IgM control (n=6), red: low-affinity anti-Insulin IgM (n=6), green: high-affinity anti-Insulin IgM (n=6). Scale bar: 10 μm. Green fluorescence indicates HEp2 cell binding. Images are representative of four independent experiments. **E:** Anti-dsDNA-IgM concentrations of insulin-specific IgM pulldowns as measured by ELISA (coating: calf-thymus dsDNA). IgM control (ctrl, n=3), IgM^low^ (n=3), IgM^high^ (n=3). Mean, ± SD, statistical significance was calculated using Kruskal-Wallis-test (all comparisons were n.s.). **F:** Insulin-binding affinity of human anti-insulin-IgM pulldowns determined by bio-layer interferometry (BLI). The K_d_ (dissociation constant) was calculated by using the K_a_ (association constant): 1/K_a_. Data are representative for three independent experiments performed in three dilution steps. **G:** Blood glucose levels of WT mice intravenously injected with 100 μg human insulin-specific IgM and human IgM control (black: n=5, red: n=4, green: n=4). Mean, ± SD, statistical significance was calculated using repeated measure ANOVA test. **H, I:** Blood glucose levels of WT mice intravenously injected with 100 μg human insulin-specific IgM (uppercase refers to affinity fraction) and human IgM control together with 500 ng native Insulin (Insulin + IgM^high^, green: n=5; Insulin + IgM control: n=5) (**H**) or together with 100 μg human anti-Insulin-IgG (n=3/group) (**I**). Mean, ± SD, statistical significance was calculated using repeated measure ANOVA test. **J:** Ratio of insulin-specific IgM of young (< 30 years, n=6) and old (> 65 years, n=6) individuals as determined by ELISA. Insulin-specific IgM was isolated via insulin-bait columns prior to ELISA experiments. Mean, ± SD, statistical significance was calculated using Kruskal-Wallis-test.

In agreement with the high specificity, the anti-insulin IgG showed no binding to any cellular structure in indirect immunofluorescence assay (IIFA) on HEp-2 cells, which is a commonly used method for detection of anti-nuclear antibodies (Buchner *et al*, 2014) (Suppl. Fig. 7). The anti-insulin IgM however, consisted of two fractions that can be biochemically separated according to their affinity to insulin (Issaq *et al*, 2002). Low-affinity anti-insulin IgM (IgM^low^) is eluted from the insulin column at higher pH (5) as compared to high-affinity anti-insulin IgM (IgM^high^) which requires acidic conditions (pH= 2.8) for elution (Fig. 2C). The IgM^low^ fraction shows polyreactivity as detected by binding to nuclear structures in IIFA and dsDNA binding in ELISA, whereas the IgM^high^ fraction is virtually negative in these assays (Fig. 2 D&E). Furthermore, we confirmed the difference in affinity by performing BLI assays and found that IgM^high^ and IgM^low^ possess a dissociation constant of 10^−10^ and 10^−7^, respectively (Fig. 2F). To test the effect of the different IgM fractions on glucose metabolism, we injected identical amounts of insulin-reactive IgM^high^ and IgM^low^ into WT mice. Increased blood glucose was observed within two hours after injection in the mice that received IgM^low^, whereas IgM^high^ did not significantly alter blood glucose levels (Fig. 2G). Moreover, we tested whether IgM^high^ plays a regulatory role under conditions of abnormally increased insulin concentrations that may cause hypoglycemia. To this end, we injected 0.1 μg insulin in combination with IgM^high^ or unspecific IgM isotype control. Strikingly, the presence of anti-insulin IgM^high^, but not the IgM isotype control, prevented the drastic decrease in blood glucose that occurred immediately after insulin injection (Fig. 2H). To further test the regulatory role of IgM^high^ in protecting insulin from IgG-mediated degradation, we combined the anti-insulin IgM^high^ with anti-insulin IgG purified from IVIg preparations. The data show that the anti-insulin IgM^high^ acts as PR-IgM as prevents the IgG-mediated neutralization of insulin which results in increased blood glucose levels (Fig. 2I, Suppl. Fig. 8). These data suggest that anti-insulin IgM^high^ is important for regulating glucose metabolism by protecting insulin from IgG-mediated neutralization and by binding excessive insulin thereby preventing drastic declines in insulin concentrations. The decrease in insulin-reactive IgM with age (Fig. 1A) prompted us to test whether the anti-insulin IgM^high^ or IgM^low^ is affected by this decrease. We determined the amount of anti-insulin IgM^high^ or IgM^low^ in young and old healthy donors and found that the ratio of anti-insulin IgM^high^ increases with age (Fig. 2J).

Together, these data suggest that glucose metabolism is regulated by different classes of antibodies and that anti-insulin IgM^high^ acts as PR-IgM that regulates glucose metabolism by regulating insulin homeostasis which seems to be particularly important with age.

### Induction of anti-insulin antibodies by insulin complexes

The presence of anti-insulin antibodies raises the question about potential mechanisms of an adaptive generation of autoreactive antibodies. We have recently shown that autoreactive antibodies can easily be elicited by polyvalent antigen in classical immunization experiments involving adjuvants (Amendt & Jumaa, 2021). To show that complexed autoantigen is capable of inducing autoreactive antibody responses independent of any adjuvants, we incubated insulin with a typical homobifunctional crosslinker, 1,2-Phenylene-bis-maleimide (Arora *et al*, 2016), which covalently binds to free sulfhydryl groups in proteins thereby crosslinking the protein of interest (Fig. 3A). Importantly, sulfhydryl group-containing drugs were reported to induce anti-insulin autoantibodies (Hu & Chen, 2018; Uchigata *et al*, 2010, 1994). Moreover, increased pancreas activity and elevated insulin production result in abnormal formation of disulfide bonds between the insulin peptides which may generate abnormal insulin forms that are more susceptible for sulfhydryl group-mediated crosslinking, and thus complex formation, under conditions of oxidative stress (Haataja *et al*, 2016; Karimi *et al*, 2016; Vinther *et al*, 2015; Van Lierop *et al*, 2017). The homobifunctional crosslinking of insulin with 1,2-Phenylene-bis-maleimide was tested in SDS page and the crosslinked insulin was purified using size exclusion spin columns excluding monomeric and dimeric insulin (Fig. 3 B). The insulin complexes were dialyzed and injected into WT mice (20 μg per mouse) without any additional adjuvants. As control, we performed a typical immunization using CpG as adjuvants and streptavidin as a foreign carrier. We found that the insulin complexes lead to increased blood glucose and anti-insulin IgM at d7 of treatment similar to the immunization (Fig. 3C, D). In addition, insulin-reactive IgG was detectable by ELISA on d14 and d26 (Suppl. Fig. 9). Repeated injection of insulin complexes at d21 resulted in further deregulation of glucose metabolism (Fig 3E). Thus, we injected anti-insulin IgM^high^ at d22, one day after injection of the insulin complexes. We found that anti-insulin IgM^high^ was able to prevent the blood glucose deregulation induced by the injection of insulin complexes (Fig. 3E).

**Figure 3.**
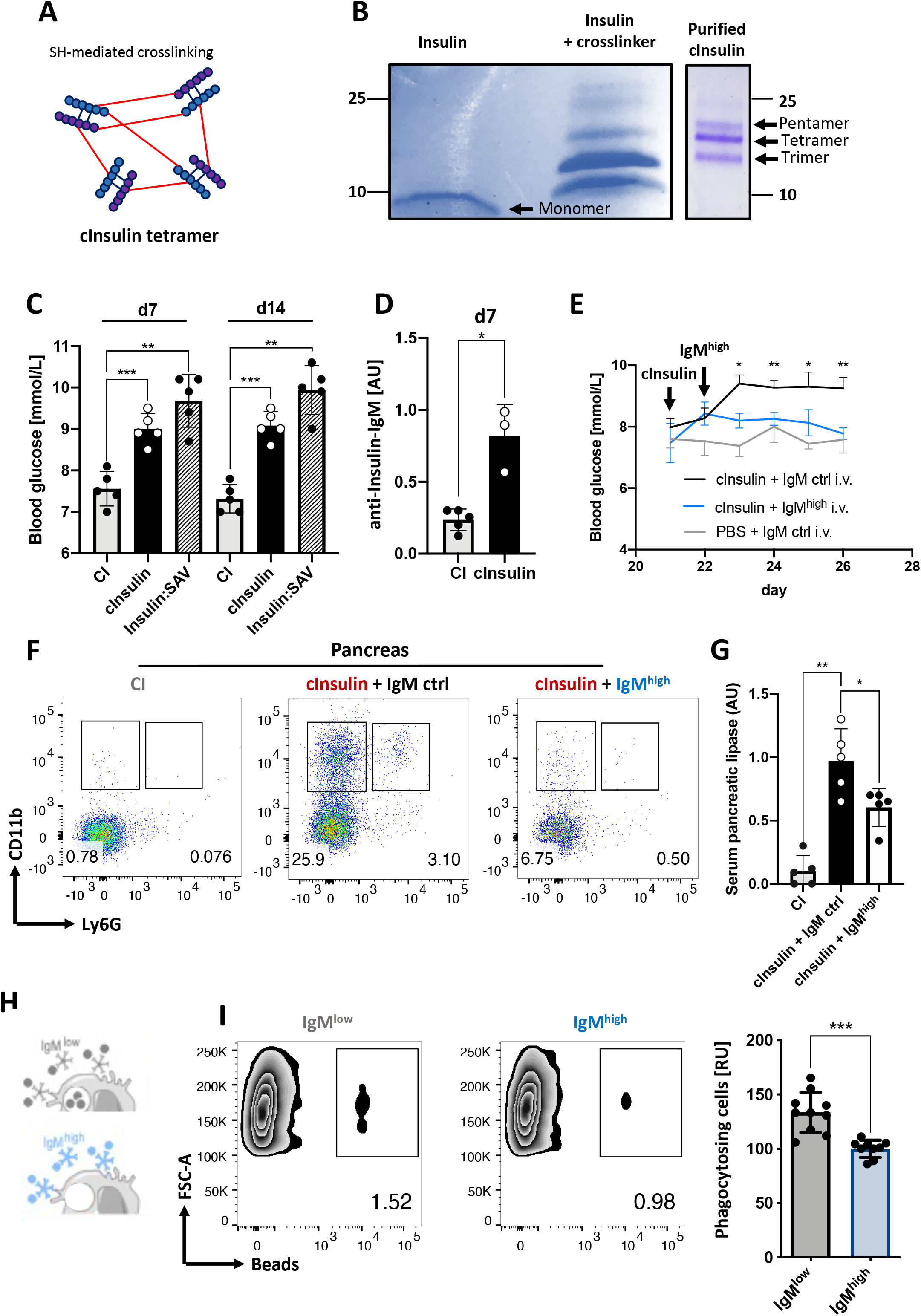
Endogenous Insulin complexes induce robust autoimmunity in mice. **A:** Schematic illustration of insulin tetramers (cInsulin) generated by thiol group-mediated disulfide bond crosslinking via 1,2-phenylene-bis-maleimide. Black lines: endogenous disulfide bonds, red lines: crosslinking induced disulfide bonds. **B:** Coomassie-stained non-reducing SDS-PAGE showing soluble Insulin (6 kD) and crosslinked insulin (> 6 kD). Right panel showing cInsulin complexes after purification with a 10 kD size exclusion column. Images are representative of three independent experiments. Marker on the left indicates band size in kilodaltons (kD). **C:** Blood glucose levels of WT mice intraperitoneally injected with PBS (control injection; CI, n=5), cInsulin (n=5) or Insulin:SAV (bio-Insulin:streptavidin, n=5) on day 0. Mean, ± SD, statistical significance was calculated using repeated measures ANOVA test. **D:** Serum anti-Insulin-IgM concentrations of WT mice intraperitoneally injected with PBS (control injection; CI, n=5) and cInsulin (n=3) on day 0 measured by ELISA at indicated days (coating: native Insulin). Mean, ± SD, statistical significance was calculated using Kruskal-Wallis-test. **E:** Blood glucose levels of WT mice intraperitoneally injected with PBS (control injection; CI, n=5) or cInsulin (n=5) on day 0 and day 21 followed by intravenous injections of 100 μg high-affinity anti-lnsulin IgM (IgM^high^) or 100 μg IgM ctrl on day 22 – 26. Mean, ± SD, statistical significance was calculated using repeated measure ANOVA test. **F:** Flow cytometric analysis of mice intraperitoneally injected with PBS (CI, n=5) or cInsulin (n=5/group) together with intravenous injections of 100 μg high-affinity anti-Insulin-IgM (IgM^high^) or 100 μg IgM control. Panels show pancreatic macrophages (CD11b^+^) and neutrophils (Ly6G^+^) pre-gated on viable cells (FVD^−^). Images are representative of three independent experiments. **G:** Serum pancreatic lipase levels of WT mice intraperitoneally injected with PBS (CI, n=5) and cInsulin (n=5/group) together with intravenous 100 μg high-affinity anti-Insulin-IgM (IgM^high^) or 100 μg IgM control. Mean, ± SD, statistical significance was calculated using Kruskal-Wallis-test. **H:** Schematic illustration of the macrophage assay used to assess phagocytosis activity of different insulin-reactive IgM affinities together with dsDNA. **I:** Flow cytometric analysis of bead-based phagocytosis assay performed with high and low affinity human anti-Insulin-IgM (IgM^low^ n=10, IgM^high^ n=10) pre-incubated with dsDNA. Left panel showing representative zebra plots and right panel showing quantitative analysis normalized to IgM^high^. Mean, ± SD, statistical significance was calculated using Kruskal-Wallis-test.

Our previous work suggests that anti-insulin IgG induces pancreas inflammation and infiltration by macrophages and granulocytes whether anti-insulin IgM^high^ was also able to prevent IgG-mediated pancreas inflammation (Amendt & Jumaa, 2021). We found that anti-insulin IgM^high^ prevents pancreas inflammation and damage as shown by the decrease of macrophage (CD11b^+^/LY6G^+^) and neutrophil (LY6G^+^) infiltration in the pancreas and the decrease of serum pancreatic lipase in blood (Fig. 3F&G).

As a mechanism for the protective role of anti-insulin IgM^high^ as compared to anti-insulin IgM^low^ we proposed that the polyreactivity of the latter, which also binds dsDNA, induces the formation of immune complexes that can be phagocytosed by macrophages, while anti-insulin IgM^high^ is highly specific for insulin and thus does not form large immune complexes that are easily phagocytosed by macrophages. To test this, we incubated anti-insulin IgM^high^ or anti-insulin IgM^low^ with insulin in the presence of genomic dsDNA (Fig. 3H). We found an increased binding/phagocytosis of anti-insulin IgM^low^ as compared with anti-insulin IgM^high^ (Fig. 3I, Suppl. Fig. 10). In addition, IgM^high^ was able to protect insulin from degradation, as the decline of insulin was greater in the supernatants containing anti-insulin IgM^low^ as compared with anti-insulin IgM^high^ antibodies (Suppl. Fig. 11).

These data show that anti-insulin antibodies can be generated under conditions activating the formation of insulin complexes, which results in deregulated glucose metabolism that can be counteracted by anti-insulin IgM^high^ that acts as PR-IgM.

### Recombinant anti-insulin IgM regulates blood glucose

The above results suggest that insulin-specific PR-IgM might be of great therapeutic interest, as it regulates insulin homeostasis and might prevent pancreas malfunction, both of which are essential for normal physiology and prevention of diabetes. According to our data, an anti-insulin IgM can act as PR-IgM if it possesses high affinity to insulin and is not reactive to autoantigens such as dsDNA or nuclear structures in IIFA. We hypothesized that a human insulin-specific IgG antibody can be converted into insulin-specific PR-IgM by exchanging the constant region.

Here, we cloned and expressed a published human insulin-specific monoclonal antibody (Ikematsu *et al*, 1994) as IgG1 (IgG_MY_) and IgM (IgM_MY_). To test the quality of our *in vitro* produced antibodies, we assessed their glycosylation by PNGaseF treatment, which resulted in reduced molecular weight as compared to untreated controls suggesting a functional glycosylation (Suppl. Fig. 12). We determined the affinity of both IgG and IgM to be 10^−9^ (K_d_) (Fig. 4A). Almost no dsDNA binding was observed in ELISA and no nuclear staining was observed in IIFA as compared to total human serum IgM (Fig 4B&C). Moreover, by injection into mice we tested if the monomeric anti-Insulin-IgM is capable of protecting insulin from degradation. Anti-Insulin IgG led to blood glucose increase which was abolished when monomeric IgM_MY_ was present (Fig. 4D).

**Figure 4.**
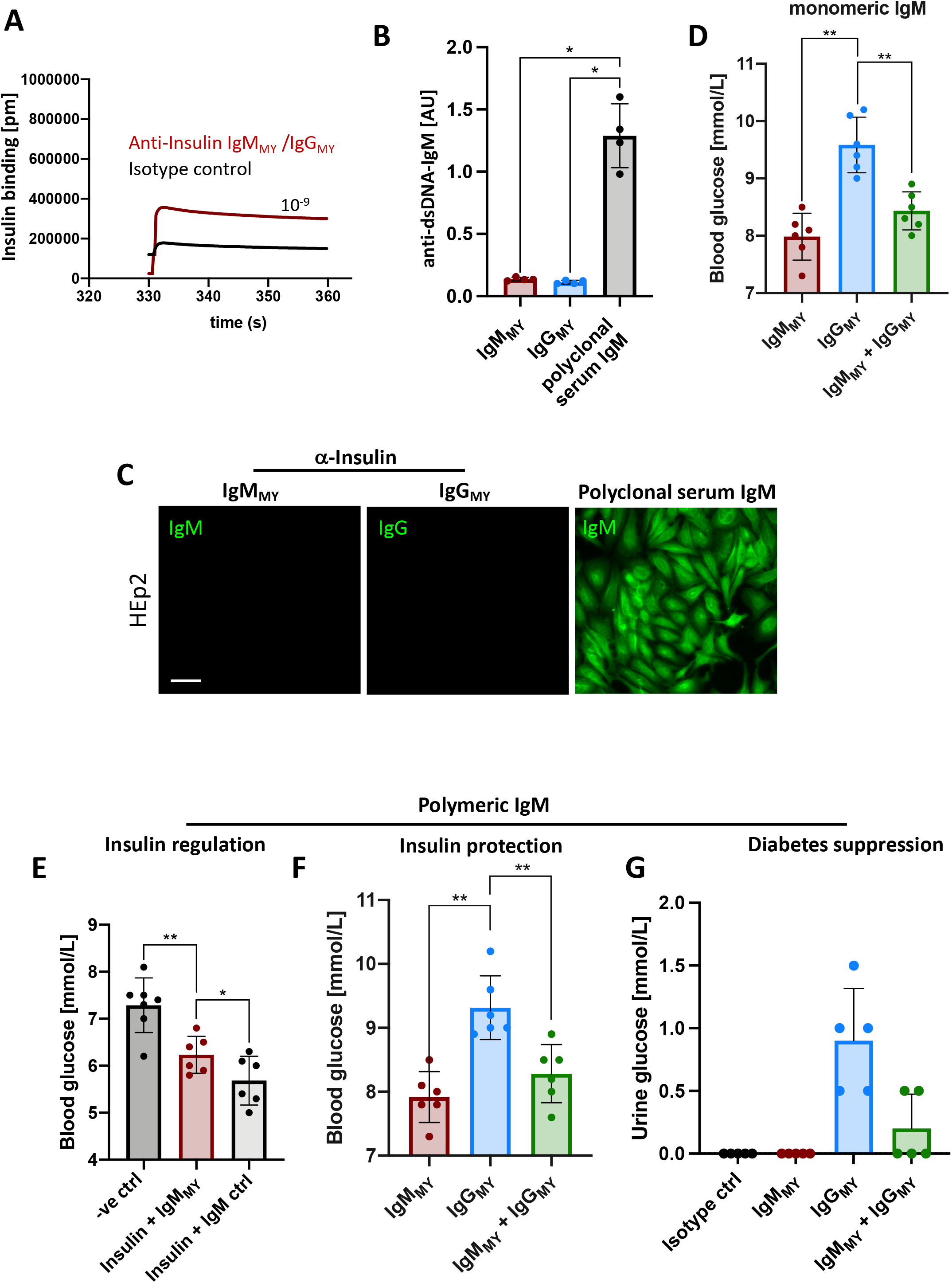
Monoclonal human insulin-IgM is able to protect Insulin *in vivo*. **A:** Insulin-binding affinity of monoclonal human anti-insulin-Ig determined by bio-layer interferometry (BLI). The K_d_ (dissociation constant) was calculated by using the Ka (association constant): 1/K_a_. Shown data are representative for three independent experiments performed in three dilution steps. **B:** Anti-dsDNA-IgM concentration of insulin-specific IgM pulldowns as measured by ELISA (coating: calf-thymus DNA). IgM_MY_ (n=4), IgG_MY_ (n=4) and polyclonal serum IgM (+ve control, n=4). Mean, ± SD, statistical significance was calculated using Kruskal-Wallis-test. **C:** HEp2 slides showing anti-nuclear structure-reactive (ANA) monoclonal IgM_MY_ (n=6), IgG_MY_ (n=6) and polyclonal serum IgM (n=6). Scale bar: 10 μm. Green fluorescence indicates HEp2 cell binding. Images are representative of three independent experiments. **D:** Blood glucose levels of WT mice intravenously injected with 100 μg monomeric IgM_MY_ (n=6), IgG_MY_ (n=6) and both combined (n=6) 2 hours post injection. Mean, ± SD, statistical significance was calculated using Kruskal-Wallis test. **E:** Blood glucose levels of WT mice intravenously injected with PBS (-ve ctrl, n=7), 500 ng Insulin with 100 μg polymeric IgM_MY_ (n=6) and 500 ng Insulin with IgM control (n=6) 2 hours post injection. Mean, ± SD, statistical significance was calculated using Kruskal-Wallis test. **F:** Blood glucose levels of WT mice intravenously injected with 100 μg polymeric IgM_MY_ (n=6), IgG_MY_ (n=6) and both combined (n=6) 2 hours post injection. Mean, ± SD, statistical significance was calculated using Kruskal-Wallis test. **G:** Urine glucose levels of WT mice immunized with cInsulin and intravenously injected with 100 μg IgM control (pentameric; n=6), 100 μg polymeric IgM_MY_ (n=5), 100 μg IgG_MY_ (n=5) and both combined (n=5) every 24 hours. Measurement shows 8 days post injection. Mean, ± SD, statistical significance was calculated using Kruskal-Wallis test.

To test whether the resulting recombinant human anti-insulin IgM_MY_ possesses protective regulatory functions, we co-injected it with insulin and found that anti-insulin IgM_MY_ prevents a drastic drop in glucose concentration induced by excess of insulin (Fig. 4E). Moreover, anti-insulin IgM_MY_ protects insulin from anti-insulin IgG_MY_ mediated neutralization, as it prevents the increase in blood glucose induced by anti-insulin IgG_MY_ (Fig. 4F). In addition, anti-insulin IgM_MY_ counteracts the leak of glucose into urine (Fig. 4G).

These data suggest that expressing a high affinity insulin-specific antibody as IgM regulates insulin homeostasis, prevents a deregulation of blood glucose concentration and might grant novel strategies for diabetes treatment.

## Discussion

This study shows that autoreactive anti-insulin IgM and IgG antibodies are normally present in wild-type animals and in human subjects. Moreover, the presence of such autoantibodies is not an accident or failure in negative selection, it is rather an important element in the regulation of insulin concentration and glucose metabolism. An even more surprising result is the finding that the affinity of autoreactive anti-insulin IgM determines the outcome of its interaction with insulin. A high affinity anti-insulin IgM is protective, whilst low-affinity IgM is destructive. Since low-affinity IgM is positive in IIFA and recognizes additional autoantigens such as dsDNA, it is multi-specific, while high-affinity IgM is monospecific, as it recognizes only insulin. The multi-specific nature of the low-affinity IgM leads to the formation of larger immune complexes (Amendt & Jumaa, 2021), which seems to accelerate insulin phagocytosis and most likely its degradation by macrophages, thereby leading to an increase in blood glucose. Obviously, the most unexpected finding is the protective role of the high-affinity IgM towards insulin, as antibody function is thought to result in neutralization of the target antigen. We refer to the generation of protective antibody as adaptive tolerance, because, in contrast to the destructive primary IgM, protective regulatory IgM is generated in the course of secondary immune responses and affinity maturation. In full agreement, our results show that anti-insulin antibody responses are induced by polyvalent insulin complexes generated by chemical crosslinking via disulfide bonds and independent of any additional adjuvants. Sulfhydryl group-containing drugs are used as immunosuppressive medication (Azathioprine(Baranzini, 2013)), anti-lymphoma treatment (Alethine(Wang *et al*, 2020)) or acute bronchitis treatment (Erdosteine(Cazzola *et al*, 2010)). Strikingly, their use was reported to result in the development of anti-insulin autoantibodies (Uchigata *et al*, 1994; Karimi *et al*, 2016). Sulfhydryl groups mediate the generation of protein complexes by the formation of disulfide bonds under oxidizing conditions (Ravasco *et al*, 2019). It is tempting to speculate that peptide hormones such as insulin or other proteins containing free sulfhydryl groups are disposed to complex formation under oxidative stress when the concentration of oxidants in the body exceeds the capacity of physiologically present antioxidants that balance oxidative stress. Thus, the anti-insulin antibody response induced in this study by oxidative crosslinking of insulin might mimic the physiological conditions which lead to autoantibody production. The oxidative stress-mediated crosslinking of insulin together with the increased amount of misfolded or abnormally crosslinked insulin peptides under conditions of elevated insulin production might lead to recurrent antibody responses against insulin. An inability of the immune system to generate PR-IgM by adaptive tolerance might represent the first step in the development of metabolic disorders associated with blood glucose. According to this scenario, type 1 diabetes develops when anti-insulin IgG production exceeds or replaces PR-IgM production resulting in failure to protect insulin and to prevent pancreas damage. On the other hand, in type 2 diabetes the production of PR-IgM may decline due to decreasing generation of new B cells with age (Frasca & Blomberg, 2011; Frasca *et al*, 2010). In full agreement, the ratio of high affinity anti-insulin IgM is highly increased in healthy aged humans as compared to young individuals. Thus, a broad B cell repertoire seems to be important for controlling autoimmunity since high-affinity protective IgM has to be produced against virtually every self-structure in order to protect it from self-destructive low-affinity IgM or IgG. This might explain the common relationship between immunodeficiency and autoimmunity (Bussone & Mouthon, 2009).

Another important conclusion of our study is that antibodies are not only required for protection from pathogens. Antibodies can also regulate the amount of key physiological factors in the blood such as insulin and are thus important regulators of metabolism and physiology. In addition to the production rate and consumption by target cells, the insulin concentration seems to be regulated by the equilibrium of protective and destructive antibodies. In this scenario, protective IgM acts as reservoir that makes insulin quickly accessible while anti-insulin IgG or low-affinity IgM neutralize the excess of insulin.

This view is supported by studies reporting severe metabolic disorders in immunodeficiency patients (Fischer, 2004). Indeed, severe combined immunodeficiency (SCID) patients with drastic hypoglobulinemia showed signs of dysglycemia and defects in fatty acid metabolism (Fischer, 2004; Webster, 1982). In full agreement, our study shows that antibody-deficient mice suffer from dysglycemia which was curable by injecting insulin-reactive autoantibodies.

Most likely, other peptide hormones or self-structures are similar to insulin regulated by autoantibodies. Methods for immortalization of memory B cells can be used for the characterization of protective IgM antibodies (Traggiai *et al*, 2004). Alternatively, converting an autoantigen-specific IgG into IgM-class antibodies might open new venues for the treatment of human autoimmune disorders by protecting the autoantigen. This might create an opportunity to turn ongoing autoimmune diseases into remission. In full agreement, we show that by converting a previously described anti-insulin IgG into IgM reversed its function from an insulin neutralizing into a protecting agent.

A further conclusion of our findings is that the highly autoreactive primary IgM repertoire represents a high risk for autoreactive damage if high affinity PR-IgM cannot be generated by secondary immune responses and somatic hypermutation. This suggests that the memory IgM repertoire consists mostly of PR-IgM generated in the course of adaptive tolerance. This also suggests that defects in somatic hypermutation leads to failure in PR-IgM generation and autoimmune damage induced by the primary IgM. In full agreement, all forms of hyper IgM syndrome (HIGM) are associated with severe autoimmunity (Durandy, 2002; Hervé *et al*, 2007; Arason *et al*, 2010; Bussone & Mouthon, 2009). HIGM patients are particularly prone to developing IgM-mediated autoimmune diseases such as immune thrombocytopenia, hemolytic anemia and nephritis (Hervé *et al*, 2007; Barbouche *et al*, 2018; Winkelstein *et al*, 2003). Notably, autoreactive IgM antibodies that cause autoimmune diseases in HIGM patients are always unmutated and therefore of low-affinity to autoantigens (Barbouche *et al*, 2018). These studies report an excessive pathogenic potential of low-affinity autoreactive IgM leading to manifest and severe autoimmunity supporting our concept of neutralizing (low-affinity) and protective (high-affinity) autoreactive IgM.

Together, our findings demonstrate an important role for autoantibodies in physiological homeostasis and suggest that adaptive tolerance is a key mechanism for maintaining physiological integrity.

## Materials and Methods

### Mice

8 – 15-weeks-old female C57BL/6 mice and mb1-deficient mice (Pelanda *et al*, 2002) were intraperitoneally (i.p.) injected with a mixture of 10 – 100 μg antigen (cInsulin or bio-lnsulin:streptavidin) in 1x PBS. Control injection (CI) mice received PBS in a total volume of 100 μL/mouse. Animal experiments were performed in compliance with license 1484 for animal testing at the responsible regional board Tübingen, Germany. All mice used in this study were either bred and housed within the animal facility of the University of Ulm under specific-pathogen-free conditions, or obtained from Charles River at 6 weeks of age. All animal experiments were performed in compliance with the guidelines of the German law and were approved by the Animal Care and Committees of Ulm University and the local government.

### Crosslinking of native Insulin

Native human insulin (Merck) was pre-diluted in water to 1 mg/mL. Chemical thiol-crosslinking was done using 1,2-Phenylen-bis-maleimide (Santa Cruz, 13118-04-2) at 100 μg/mL for 48 h at room temperature (dark) and afterwards removed by using a 10 kD cut-off spin column (Abcam, ab93349). Purified insulin complexes (cInsulin) were used for intraperitoneal injections. Quality control was done by using a non-reducing SDS-PAGE and Coomassie staining (see below).

### Macrophage phagocytosis assay

The effect of IgM^high^ and IgM^low^ on the antibody-mediated clearance was assessed in mouse phagocytic J774A.1 cell line. Red fluorescent latex beads (2 μm in diameter, Sigma-Aldrich) were coated with purified IgM^high^ and IgM^low^ in a 1:7 ratio and incubated for 45 min at 37°C IgM Promotes the Clearance of Small Particles and Apoptotic Microparticles by Macrophages (Litvack *et al*, 2011). The IgM-coated latex beads were suspended in serum-free culture medium and added to J774A. 1macrophages at a 5:1 ratio. After 180 min incubation, cells were washed in PBS to remove non-engulfed beads. Cells were detached with trypsinization and fixed in 4% PFA, and subsequently analyzed with BD LSR II flow cytometer. The macrophage population was identified as described before (Yu *et al*, 2019) and the proportion of macrophages containing ingested particles was determined based on the fluorescent signal of the ingested latex beads.

### Flow cytometry

Detailed description is available in Setz *et al*, 2019. Briefly, cell suspensions were Fc-receptor blocked with polyclonal rat IgG-UNLB (2,4G2; BD) and stained according to standard protocols. Viable cells were distinguished from dead cells by using Fixable Viability Dye eFluor780 (eBioscience) and stained cells were acquired at a Canto II Flow Cytometer (BD).

### Enzyme-linked Immunosorbent Assay (ELISA)

Antigen-specific ELISA assays are described in Amendt & Jumaa, 2021 in detail. Briefly, 96-Well plates (Nunc, Maxisorp) were coated with, native Insulin (Sigma-Aldrich, Cat. 91077C), calf thymus dsDNA (ThermoScientific, Cat.15633019), at 10 μg/mL. Standard coating was done by using anti-IgM, anti-IgG-antibodies (SouthernBiotech). The relative concentrations, stated as arbitrary unit (AU), or absolute concentration (μg/mL), were determined via detection by alkaline phosphatase (AP)-labeled anti-IgM/anti-IgG (SouthernBiotech) by using p-nitro-phenylphosphate (pNPP; Genaxxon) in diethanolamine buffer, respectively.

### Enzyme-linked Immuno-Spot Assay (ELISpot)

Total splenocytes were measured in triplicates with cell numbers stated in the figure. ELISpot plates were coated with native Insulin (Sigma-Aldrich, Cat. 91077C), anti-IgM (Mabtech) or anti-IgG (Mabtech). Seeded cells were incubated for 12 – 24 h at 37 °C, antigen-specific IgM or IgG was detected via anti-IgM-bio and SAV-AP or anti-IgG-bio and SAV-AP (Mabtech). Experiments were performed according to the manufacturer’s instructions.

### HEp-2 slides and fluorescence microscopy

HEp2 assays with corresponding autoantibodies are described in Amendt & Jumaa, 2021 in detail. Briefly, serum samples indicated in the figures or figure legends were diluted to an equal concentration of IgM (approx. 300 ng/mL anti-Insulin-IgM in both immunized samples) and applied onto the HEp-2 slides (EUROIMMUN, F191108VA). Anti-IgM-FITC and anti-IgG-APC (eBioscience) were used for visualization of ANA-IgM/-IgG.

### Glucose level monitoring

Assessment of blood and urine glucose levels is described in Amendt & Jumaa, 2021 in detail. Briefly, we used an AccuCheck (Roche Diagnostics, Mannheim) blood glucose monitor (mmol/L) and took blood from the tail vein.

### SDS-PAGE

Samples were separated on 10 – 20 % SDS-polyacrylamide gels which were incubated with Coomassie (Coomassie brilliant blue R-250, ThermoFisher) for 45 min and subsequently de-stained.

### Pulldown of total serum immunoglobulins

Serum of injected mice was collected immediately after euthanization at indicated days. Antigen bound to antibodies was removed by repeated freeze-thaw cycles of the serum and pH-shift during elution (Reverberi & Reverberi, 2007). Protein G sepharose beads (ThermoFisher) were used according to the manufacturers protocol and obtained antibodies dialyzed overnight in 10 times sample volume of 1x PBS. For isolation of IgM, HiTrap IgM columns (GE Healthcare, Sigma-Aldrich) were used according to the manufacturers protocol and dialyzed overnight in 10 times sample volume of 1 x PBS. The quality of the isolated immunoglobulins was checked via SDS page and Coomassie-staining and the concentration of antigen-specific immunoglobulins determined by ELISA.

### Isolation of antigen-specific immunoglobulins

Streptavidin bead columns (ThermoScientific, Cat. 21115) were loaded with 10 μg bio-Insulin (BioEagle) in order to isolate insulin-specific immunoglobulins. Samples (sera or isolated antibodies) were incubated for 90 min at room temperature to ensure sufficient binding of antibodies to the insulin-loaded beads. Purification of the insulin-antibodies was achieved by a pH-shift (pH 2.8) using the manufacturers elution and neutralization solutions. Note, fractionation into low and high affinity fractions was done by eluting with pH 5 (low affinity) and subsequently with pH 2.8 (high affinity). The quality of the isolated immunoglobulins was determined via reduced and non-reduced Coomassie-stained SDS-PAGEs and the quantity determined by ELISA. For further *in vivo* experiments, the isolated antibodies were dialyzed as described above.

### Bio-Layer-Interferometry (BLI)

Interferometric assays (BLItz device, ForteBio) were used to determine the affinity of protein-protein interactions (Kumaraswamy & Tobias, 2015) as described in Amendt & Jumaa, 2021.

### Wire hanging test

The linear wire hanging test was used to assess motor strength and function of mice(Hoffman & Winder, 2016). Individual mice were put onto a 36 cm elevated horizontal wire above a cage, subsequently the mice tried to stay on the wire by using their paws and muscle strength. The ability in time (sec) of each mouse to stay on the wire was recorded. A maximum time duration of 240 sec was set. Each mouse went through the test three times in a row. The mean value was calculated from the measured data. Blood glucose values were determined before and after the test.

### Rotarod test

To assess motor function and blood glucose levels after physical exercise, the rotarod test was performed(Hamm *et al*, 1994). Before the start of the experiment, the blood glucose level of the mice was determined. Afterwards they were placed on the running drum of the Rotarod apparatus with a continuous rotation of 32 rpm. At 30 sec the experiment was stopped and the blood glucose values were measured.

### Statistical analysis

Graphs were created and statistical analysis was performed by using GraphPad Prism (version 6.0h) software. The numbers of individual replicates or mice (n) are stated within the figure or figure legends. Data sets were analyzed by D’Agostino & Pearson omnibus normality test and/or Shapiro-Wilk normality test in GraphPad Prism software to determine whether they are normally distributed. If one of the data sets was not normally distributed or the sample number n was too small to perform the normality tests, non-parametric tests were used to calculate p-values. In this study, p-values were calculated by tests stated in the respective figure legends. Students t-tests with Welch’s correction were used to compare two groups within one experiment. P values > 0.05 were considered to be statistically significant (n.s.=not significant; * p < 0.05; ** p < 0.01; *** p < 0.001, **** p < 0.0001).

## Author contributions

TA performed experiments, analyzed and interpreted data, prepared the figures and contributed to the writing of the manuscript. GA performed behavior tests with mice. GA, AN, MY, OEA and CSS performed ELISA experiments. HY and TR designed and performed macrophage phagocytosis assays. MR and KMD helped with discussions on the manuscript. HJ designed the study, proposed experiments, wrote the manuscript and supervised the project.

## Acknowledgements

We thank the group of T. Böcker for the free use of the mouse behavior experimental room. M. Reth for critical reading and fruitful discussion. This work was supported by the DFG through TRR130 (B cells and beyond) project 01, SFB1074 (Experimental Models and Clinical Translation in Leukemia), SFB 1279 (Exploration of the Human Peptidome), and DFG single grant, JU 463/5-1.

