## Supplementary Figures for "Antibodies control metabolism by regulating insulin homeostasis"

**Suppl. Fig. 1**

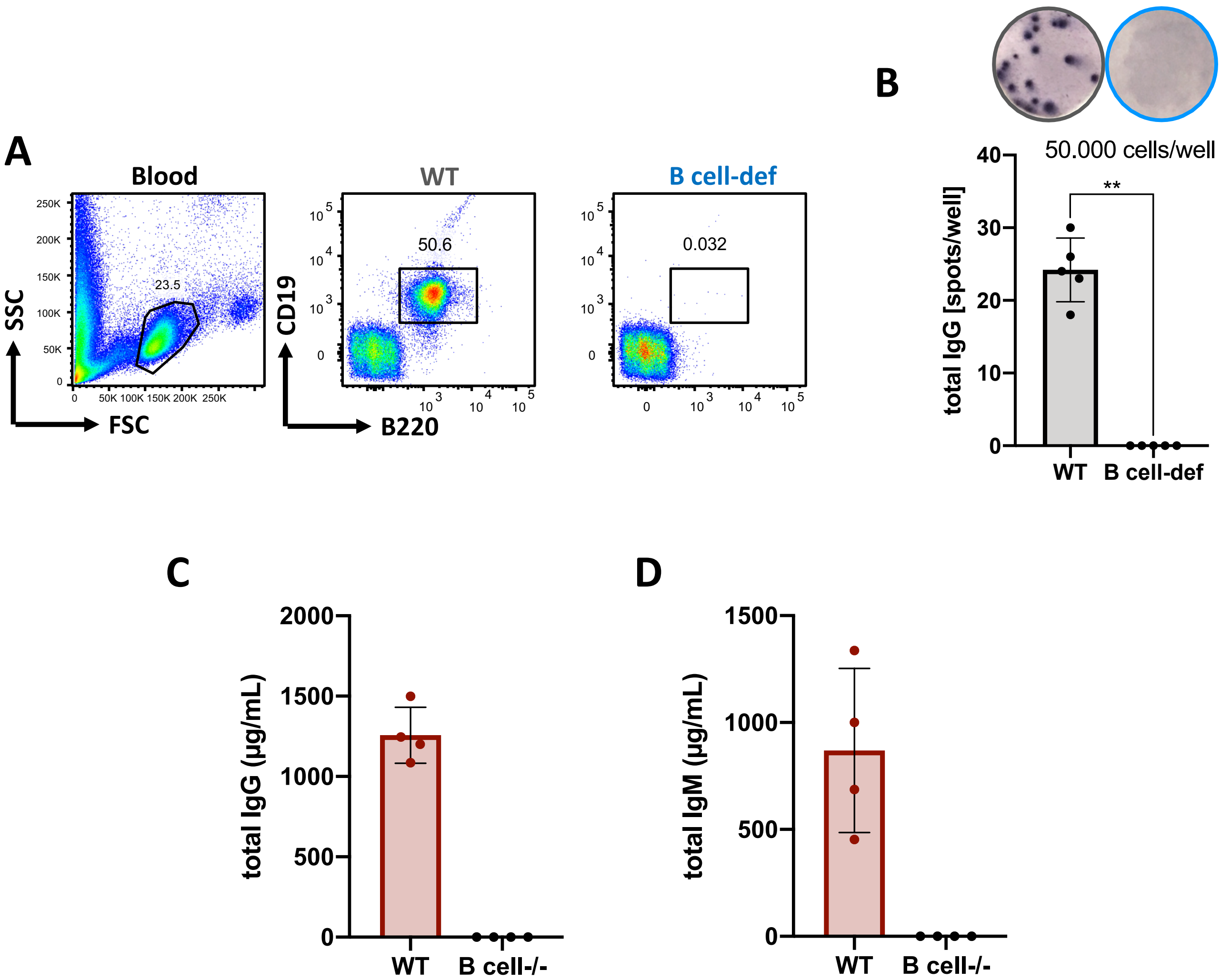


**Figure S1 | No antibody secreting cells in B cell-deficient mice.**

**A**: Flow cytometric analysis of blood of wild-type (red) and B cell-deficient (blue) mice. Left panel shows size and granularity of cells in forward and sideward scatter. Middle and right panel showing cells pre-gated on lymphocytes.

**B**: IgG secreting splenocytes of wild-type and B cell-deficient mice measured by ELISpot. 50.000 splenocytes were seeded per well. Mean ± SD, statistical significance was calculated by using Kruskal-Wallis-test.

**C, D**: Serum total IgG (C) and total IgM (D) titers of wild-type and B cell deficient mice as measured by ELISA. Mean ± SD, statistical significance was calculated by using Kruskal-Wallis-test.

**Suppl. Fig. 2**

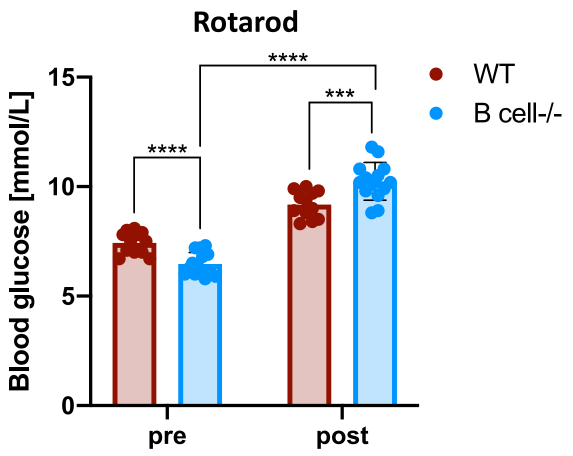


**Figure S2 | B cell-deficient mice show dysglycemia after exercise.**
Blood glucose levels of wild-type and B cell-deficient mice post rotarod exercise. Animals were placed on a rotarod for 60 sec and subsequently blood glucose levels analyzed. n=12/group, mean ± SD, statistical significance was calculated by using Kruskal-Wallis-test.

**Suppl. Fig. 3**

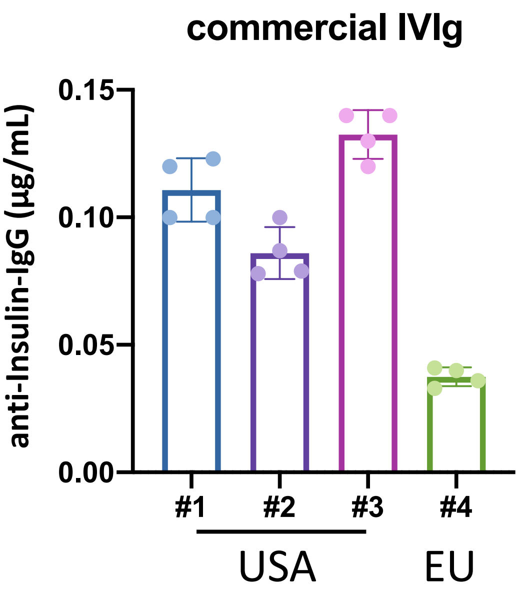


**Figure S3 | Insulin-specific IgG in commonly used IVIg preparations.**
Anti-Insulin-IgG of commercial IVIg formulations as measured by ELISA (coating: native Insulin). IVIg preparations were diluted to 1 mg/mL. n=4/group, mean ± SD, statistical significance was calculated by using Kruskal-Wallis-test.

### Suppl. Fig. 4

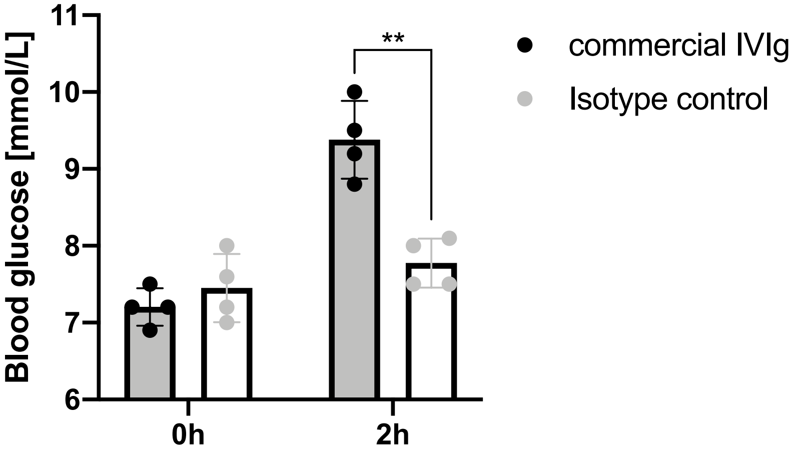


**Figure S4 | IVIg injections lead to increased blood glucose levels in wildtype mice.**
Blood glucose levels of wildtype mice intravenously injected with 200 µg commercial human IVIg and 200 µg isotype control (mAb IgG1, IgG2, IgG3, IgG4 mix) measured by commercial blood glucose monitoring at indicated hours. Mean ± SD, statistical significance was calculated by using repeated measure ANOVA-test.

### Suppl. Fig. 5

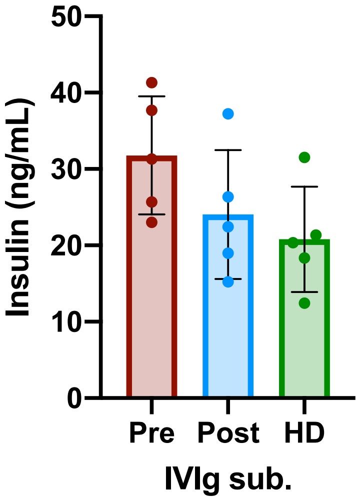


**Figure S5 | Immunodeficiency patients treated with commercial IVIg suffer from transient Insulin depletion.**
Serum insulin levels of immunodeficiency patients treated with commercial human IVIg preparations measured by ELISA. n=4/group, mean ± SD, statistical significance was calculated by using Kruskal-Wallis-test.

### Suppl. Fig. 6

**A**

| Insulin epitope | Amino acid sequence |
|---|---|
| InsA-1 (99 – 110) | CSLYQLENYCN |
| InsA-2 (88 – 98) | GIVDQCCTSI |
| InsB-1 (25 – 35) | FVKQHLCGPHL |
| InsB-2 (36 – 46) | PHLVEALYLVCG |
| InsB-3 (57 – 58) | CGERGFFYTPKS |

**B**

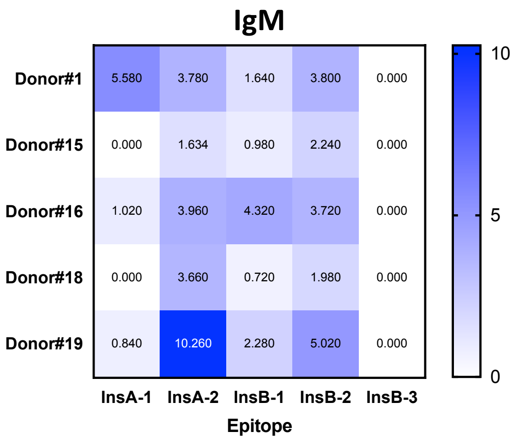


**Figure S6 | Human anti-Insulin IgM is not restricted to a single epitope.**
**A:** Table showing sequences for Insulin-A and B chain derived epitopes used as peptides.
**B:** Serum anti-Insulin-IgM of human serum samples as measured by ELISA. n=5, mean ± SD, statistical significance was calculated by using Kruskal-Wallis-test.

### Suppl. Fig. 7

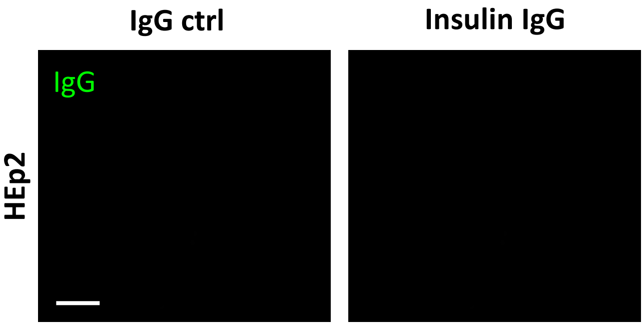


**Figure S7 | Human anti-Insulin IgG is present in healthy individuals and is monospecific.**
HEp2 slides of human IgG recognizing Insulin and IgG control showing nuclear structure binding antibodies. Scale bar: 10 µm.

#### Suppl. Fig. 8

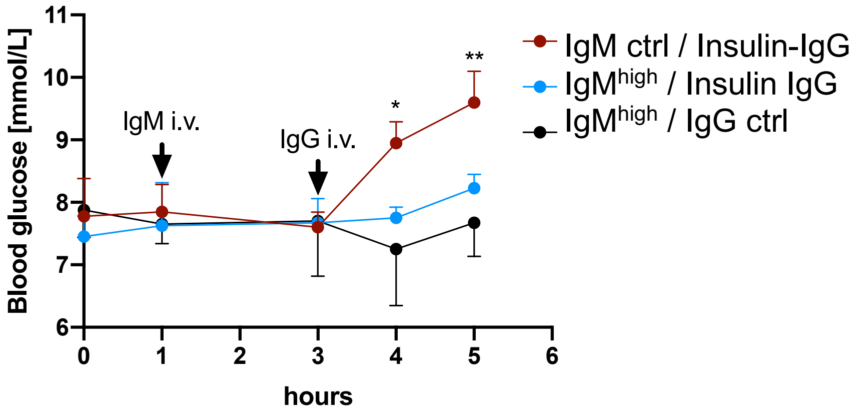


**Figure S8 | Protective IgM is able to suppress hyperglycemia when competing with anti-insulin IgG.**
Blood glucose values of WT mice intravenously injected with anti-Insuiln high affinity IgM (IgM$^{high}$) and anti-Insulin IgG. n=6/group, mean ± SD, statistical significance was calculated by using repeated measure ANOVA.

### Suppl. Fig. 9

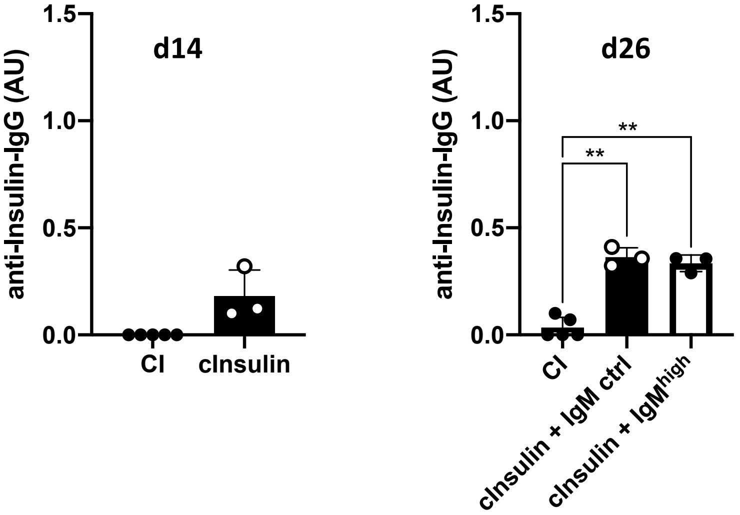


**Figure S9 | Injections with cInsulin induce insulin-reactive IgG and injections with high-affinity IgM prevents diabetes without blocking the immune reaction.**
Serum anti-Insulin-IgG of cInsulin and control injected (PBS) mice measured at indicated days by ELISA. n=3/group, n=5 for controls, mean ± SD, statistical significance was calculated by using Kruskal-Wallis-test.

### Suppl. Fig. 10

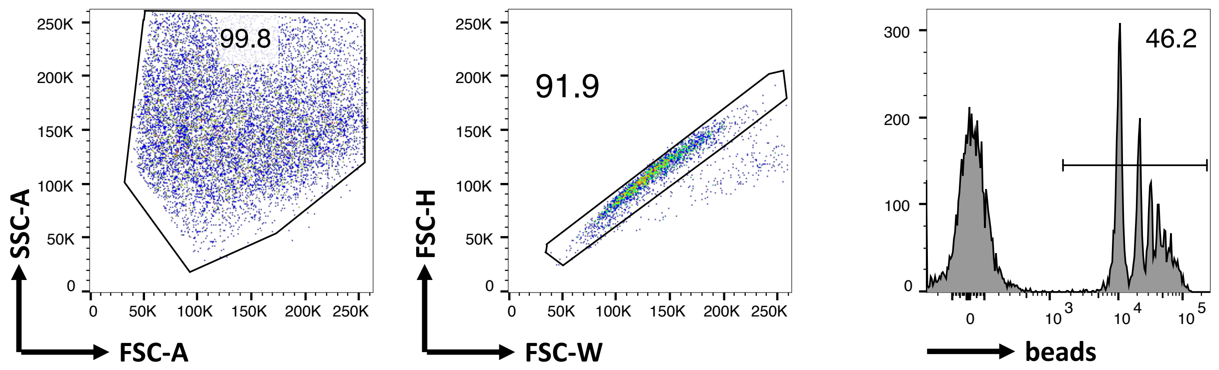


**Figure S9 | Gating strategy and experimental set-up for macrophage phagocytosis assays.** Flow cytometric analysis of macrophage bead phagocytosing assay. Left panel showing macrophages in SSC-A and FSC-A used for bead phagocytosis. Middle panel showing single cells pre-gated on macrophages. Right panel showing phagocytosing macrophages (beads$^{+}$) and non-phagocytosing cells (beads$^{-}$). Images representative of two independent experiments.

**Suppl. Fig. 11**

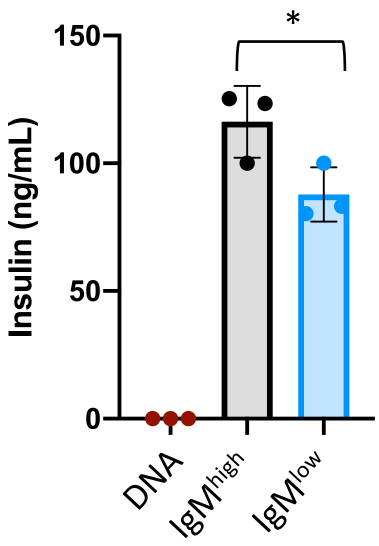


**Figure S10 | IgM$^{high}$ protects Insulin from macrophage-mediated neutralization.**
ELISA detecting Insulin levels of macrophages treated with immune complexes of DNA + Insulin + either low or high affinity IgM. n=3/group, mean ± SD, statistical significance was calculated by using Kruskal-Wallis-test.

### Suppl. Fig. 12

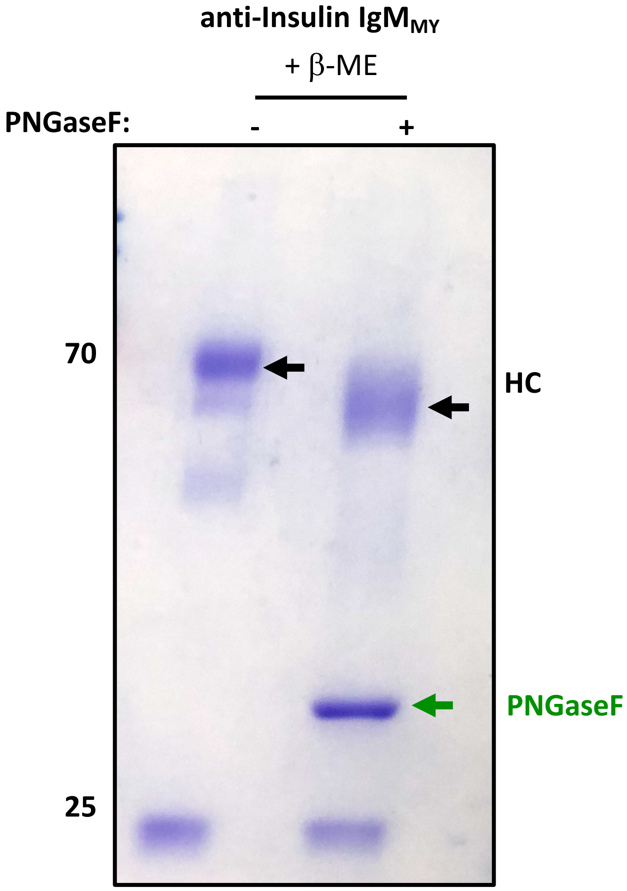


**Figure S11 | Monoclonal anti-Insulin IgM is physiologically glycosylated.**
Coomassie stained SDS-page of IgM pentamers treated with PNGaseF. Beta-Mercaptoethanol was used to reduce the antibodies. Image representative of two independent experiments. Black arrows mark the IgM heavy chain (HC). Green arrow marks PNGaseF enzyme.
